# Semantic object processing is modulated by prior scene context

**DOI:** 10.1101/2022.10.26.513851

**Authors:** Alexandra Krugliak, Dejan Draschkow, Melissa L.-H. Võ, Alex Clarke

## Abstract

Objects that are congruent with a scene are recognised more efficiently than objects that are incongruent. Further, semantic integration of incongruent objects elicits a stronger N300/N400 EEG component. Yet, the time course and mechanisms of how contextual information supports access to semantic object information is unclear. We used computational modelling and EEG to test how context influences semantic object processing. Using representational similarity analysis, we established that EEG patterns dissociated between objects in congruent or incongruent scenes from around 300 ms. By modelling semantic processing of objects using independently normed properties, we confirm that the onset of semantic processing of both congruent and incongruent objects is similar (∼150 ms). Critically, after ∼275 ms, we discover a difference in the duration of semantic integration, lasting longer for incongruent compared to congruent objects. These results constrain our understanding of how contextual information supports access to semantic object information.

## Introduction

In our daily lives, we easily recognise the objects around us. Yet, in certain situations we expect to see some objects more than others. For example, when walking down a street, we might expect to encounter a car but not an elephant. But if we visit a zoo, we would be much more likely to encounter an elephant in an enclosure than a car. In both scenarios, we recognise the object as a car and an elephant, however, the context in which we see these objects influences the way we perceive and respond to them. Objects that are congruent with their environment are recognised faster and more accurately than objects that are incongruent (Bar, 2004; Biederman et al., 1982; Davenport & Potter, 2004; Greene et al., 2015; Oliva & Torralba, 2007; Palmer, 1975). This is also reflected in neural processing, in that incongruent objects induce a stronger negativity of the N300/N400 EEG components than congruent objects (e.g. Draschkow et al., 2018; Ganis & Kutas, 2003; Lauer et al., 2018; Lauer et al., 2020; Mudrik et al., 2010; Mudrik et al., 2014; Võ & Wolfe, 2013). Such congruency effects for stimuli mismatching a context have been reported not only for objects but for a variety of stimuli, like words at the end of a sentence (e.g. Kutas & Hillyard, 1980), images in a preceding sentence context (e.g. Ganis et al., 1996), or scene images preceded by a verbal cue of a scene category (Kumar et al., 2021), indicating that during the N300/N400 interval, semantic information becomes available and is integrated into the context (for a review see Kutas & Federmeier, 2011).

Much of what we do know about the semantic processing of visual objects comes from research where objects are presented isolated from the background or in a stream of unconnected events. This line of research indicates that in the first ∼150 ms after the object appears, low- and middle-level object features are extracted, mostly in a feedforward fashion along the ventral visual stream (Cichy et al., 2016; DiCarlo et al., 2012; Lamme & Roelfsema, 2000). More complex visual features and semantic features are processed at later latencies, beginning after 150-200 ms, supported by recurrent dynamics within the ventral temporal lobes (Bankson et al., 2018; Chan et al., 2011; Clarke, 2019; Clarke et al., 2011, 2018; Kietzmann et al., 2019; Poch et al., 2015). In agreement with this object processing timeline, the effects of object-scene congruency on the N300/N400 EEG components occur at a similar time as semantic feature effects for single objects, which would allow for context to modulate object perception.

Previous EEG research suggests the scene context can directly modulate the timing of object processing. For instance, Truman and Mudrik (2018) reported that when intact and scrambled objects were shown embedded in congruent or incongruent scenes, EEG signals to intact objects in congruent contexts diverged from EEG signals to scrambled images within the N300 time window, while EEG signals to intact objects in incongruent contexts diverged in the N400 time window. They suggest that object identification is delayed when objects are in incongruent scenes and integration of the object and scene is enhanced. While this might suggest differences in the timing and duration of semantic access for objects in congruent and incongruent scenes, the research so far is limited in answering the question of how semantic object information is represented in these different situations, in terms of the timing of semantic activation and the nature of this semantic information. Contrasts between congruent and incongruent conditions are well suited to exploring differences in processing between these conditions, while understanding how we access semantic information for objects in different contexts is better aided through approaches that measure semantic processing individually for each of these conditions. By tracking the semantic processing of objects in congruent scenes, separately from the semantic processing of objects in incongruent scenes, we can more directly test for differences and similarities in how semantic information is accessed.

Three plausible scenarios for how semantic access is modulated by a prior scene context are (1) that semantic object information is accessed faster for objects in congruent compared to incongruent environments, meaning that later processing of objects in incongruent environments leads to a N300/N400 congruency effect, (2) semantic access is initiated at the same time in both conditions, but continues for longer in the incongruent case, with the additional semantic activation related to congruency effects, or (3) that semantic access is initiated at the same time and for the same duration for both congruent and incongruent conditions, and differences in the magnitude of semantic access relate to congruency effects.

Here, we re-analysed EEG data by Draschkow and colleagues (2018) to test the hypothesis that neural effects of congruency on the N300/N400 components are driven by differences in accessing semantic object information, by combining computational models with Representational Similarity Analysis (RSA; Kriegeskorte et al., 2008) - a methodology that allows testing specific hypotheses about what object features contribute to neural signals during object processing (Bankson et al., 2018; Cichy et al., 2016; Clarke et al., 2018). During RSA, the similarity of neural responses between individual objects is calculated and summarized in a Representational Dissimilarity Matrix (RDM). These RDMs of brain signals can be calculated at each point in time, and tested against a second set of RDMs that represent our predictions for why objects might be more or less similar to one another (e.g. due to congruency or semantic similarity). A significant relationship between the neural RDMs and RDMs of our predictions (or models), suggests that the predicted information is currently being represented in neural signals. For example, Clarke and colleagues (2018) demonstrated this approach using a model of semantics based on features from a property norming study (Devereux et al., 2014), which was related to MEG signals. The property norms were obtained by asking participants to name features associated with concept words, resulting in a collection of 3026 different features that capture the semantic representations of individual concepts (e.g. a car has the features ‘has wheels’, ‘has a driver’ and ‘made of metal’ but not the features ‘is edible’, ‘has wings’), which then allows an examination of the relationship between neural responses to single objects and the semantics defined by the norms. Clarke and colleagues (2018) reported the semantic model, based on such property norms, related to brain activity peaking around 250 ms after object onset - a latency similar to the onset of the N300/N400 component. Using a similar model here, based on the same independently normed semantic features as used by Clarke and colleagues (2018), provides not only the intriguing opportunity to directly test if context indeed modulates semantics, but also how it effects the temporal processing of objects in congruent and incongruent settings independently, adjudicating between the three scenarios we set out above.

In the current EEG data set, participant viewed images of scenes where a cue indicated the location where either a congruent or incongruent object would appear. We extracted similarity of neural responses to objects and related them to different models. First, we used a simple congruency model that distinguishes between congruent and incongruent contexts, to establish when representational differences in the EEG signals emerge that suggests a dissociation of processing between the conditions. Then we modelled the EEG data with a semantic model based on property norms that describes the objects in terms of semantic features, to specifically test how congruency impacts the time course of processing semantic object information, and how this is different depending on the contextual congruency between the object and the scene.

## Methods

We re-analysed EEG data reported by Draschkow and colleagues (2018). The data is freely available (https://github.com/DejanDraschkow/n3n4). Here we provide a short summary of the main aspects of the study design covering participants, procedure, EEG recording and pre-processing, as well as the specifications of our RSA analyses.

### Participants and procedure

Forty healthy participants viewed 152 scene images that were presented with either a semantically congruent or incongruent object (76 trials per condition). Each scene was paired with a congruent and an incongruent object, but participants saw each scene only once with either the congruent or the incongruent object (the conditions were counterbalanced across participants). At the beginning of each trial, a scene was presented for 500 ms, then a red dot appeared indicating the position where the object would appear. After 500 to 530 ms, the object was presented in the cued location of the scene and remained on the screen for 2000 ms (Fig 1, Fig 2A). The task was to report exact repetitions of scenes and objects (the repetition trials were excluded from subsequent analysis).

**Figure 1.**
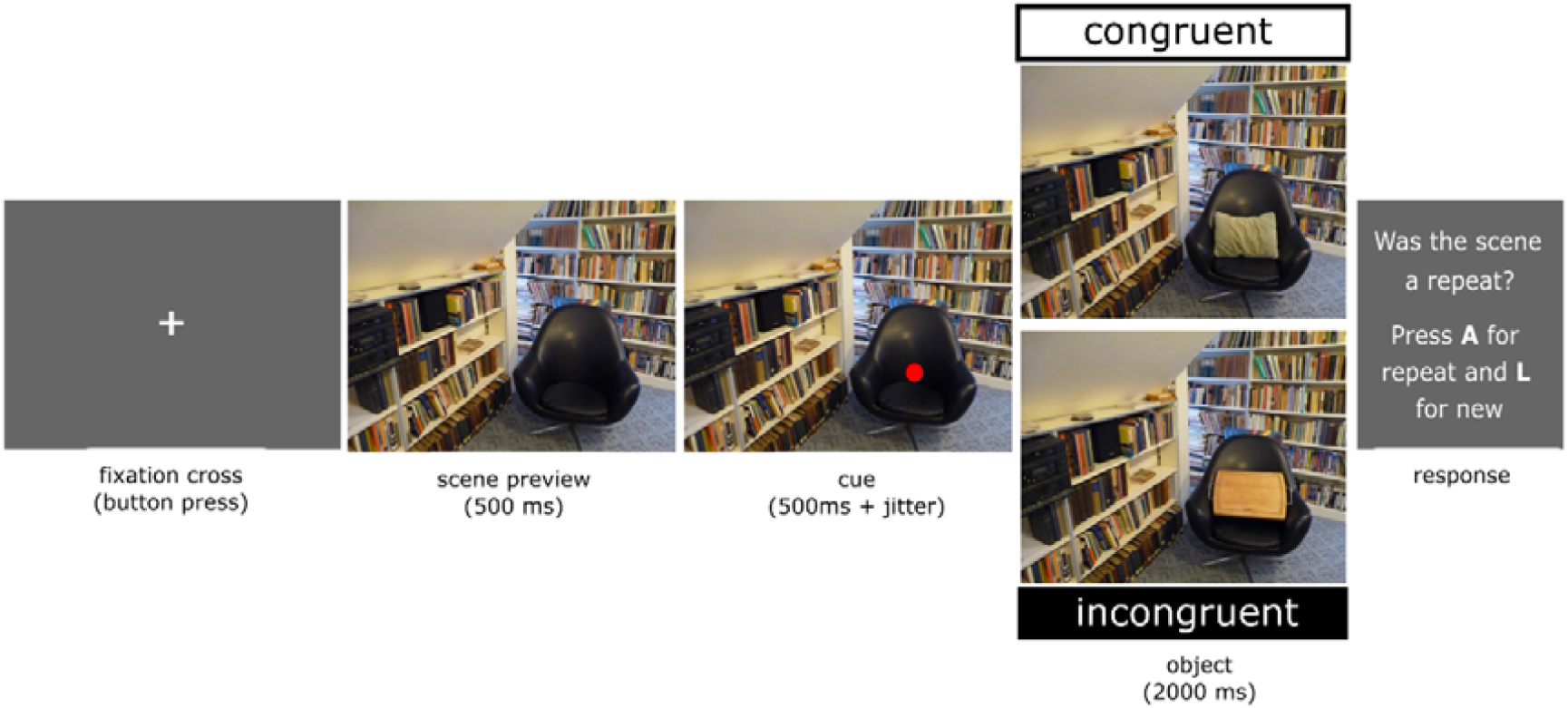
An example trial showing a scene before a red dot appears to indicate the location the object will appear. The object that appeared could either be congruent with the scene, in this example a cushion, or incongruent with the scene, in this example a chopping board. Each scene is only shown once to a participant, with either a congruent or incongruent object.

**Figure 2.**
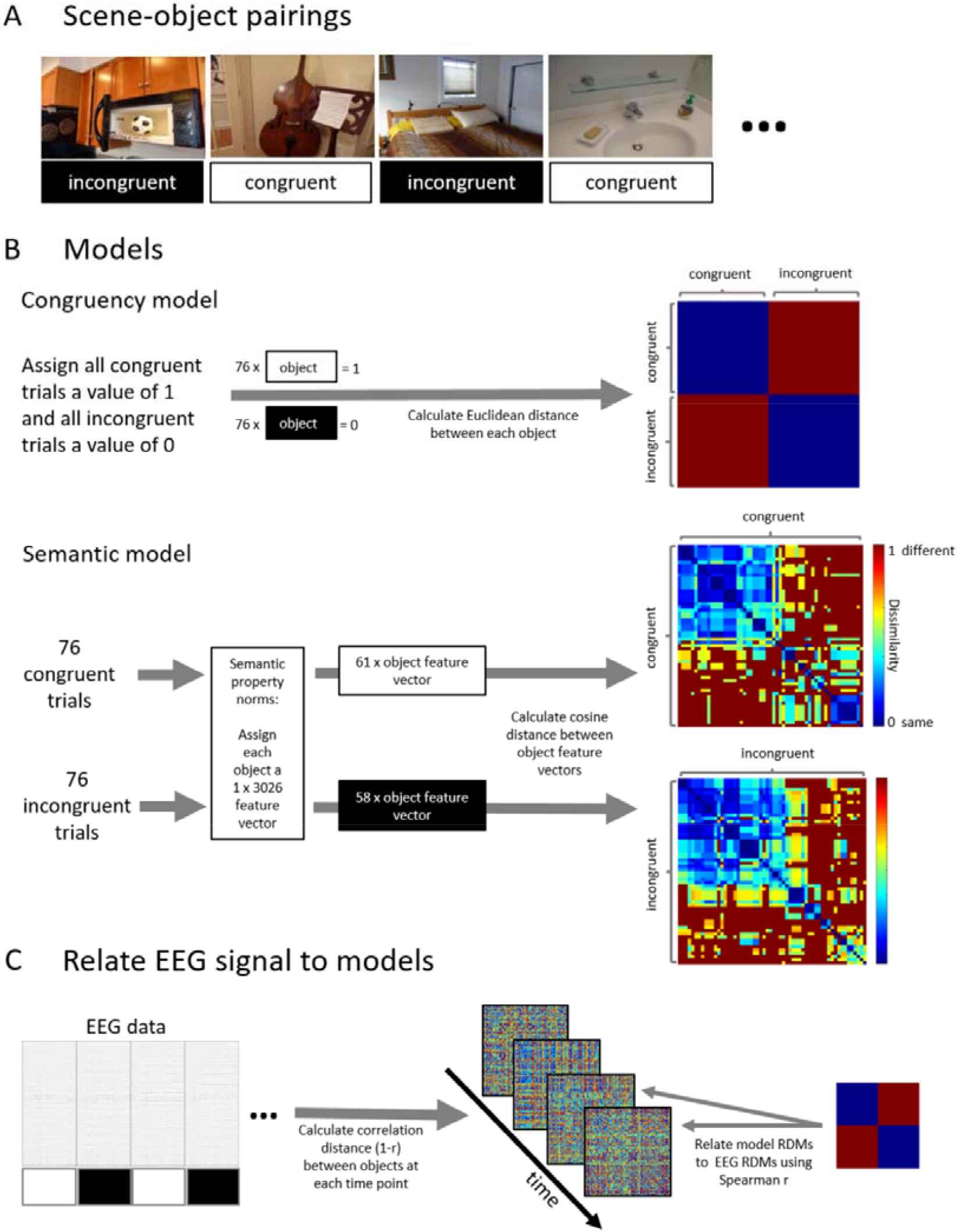
An overview of Representation Similarity Analysis (RSA) that relates brain responses at each time point to the congruency and semantic information associated with the different trials. (A) Example stimuli showing objects in congruent and incongruent scenes. (B) Model RDMs for an example participant. For the congruency model analysis all congruent object trials were each assigned a value of 1 and all incongruent object trials each a value of 0. Then the Euclidean distance between all object-pairs was calculated. The resulting RDM directly dissociates congruent and incongruent objects. For the semantic model analysis, the congruent and incongruent trials were analysed separately. Each object for which independently normed semantic features were available was assigned a corresponding semantic feature vector. Then the cosine distance was calculated between the feature vectors of all object-pairs, separately for consistent and inconsistent trials, resulting in two RDMs that describe the similarity of objects based on semantic properties. (C) The model RDMs are then statistically related to brain signals. For each object, the EEG response was extracted across all channels, and at each time-point the similarity between object pairs was calculated using correlation distance. Model RDMs were then related to these EEG RDMs using Spearman correlation.

**Figure 3.**
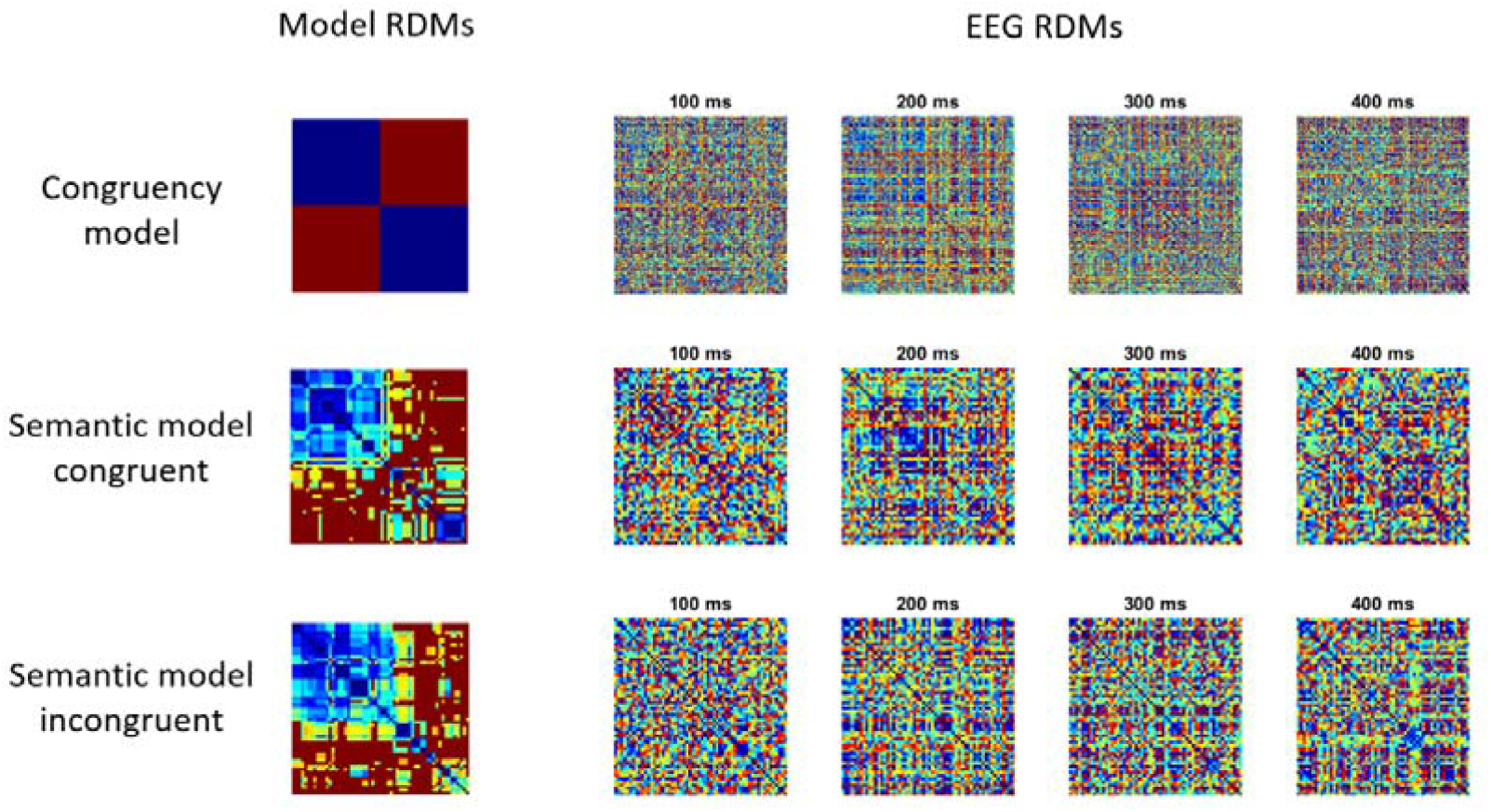
Model RDMs tested and EEG RDMs from an example participant at different time points.

### EEG recording and pre-processing

EEG data was recorded with 64 active Ag/AgCl electrodes (Brain Products, GmhB), with a sampling rate of 1000 Hz. Data were down-sampled to 200 Hz, filtered between 0.1 Hz and 40 Hz, and eye and muscle artifacts were removed with independent component analysis. Epochs of 1100ms were created, from -200 ms to 900 ms centred around object onset, then baseline correction was applied from -200 ms to 0 ms. We started our analysis with the epoched data provided by Draschkow and colleagues (2018), however, we identified noisy trials by visual inspection, specifically those trials that contained high frequency noise or large amplitude signals beyond the range of normal activity. On average 2.9% of trials were removed (range 0-24%). All electrodes were included in the subsequent Representation Similarity Analysis (RSA).

### Representational Similarity Analysis

RSA was used to relate model-based congruency and semantic similarity between objects to the neural similarity based on EEG data (Figure 2). We computed the similarity between neural responses to objects at each moment in time as 1-Pearson correlation between the EEG signals for each object pair and summarized the similarity measures in symmetric Representational Dissimilarity Matrices (RDMs) per time-point. Then, using Spearman correlation, we related the EEG RDMs to RDMs that reflect a similarity structure between objects based on either the congruency or semantic properties. This method allows to reveal when and for how long the EEG signals distinguish between objects based on the contextual congruency or the semantic properties of those objects.

#### Congruency model analysis

The congruency model dissociates between objects presented in congruent and incongruent contexts. We created participant-specific models because the same scenes were presented to some participants with a congruent object and to other participants with an incongruent object (counterbalanced across participants). For each participant, we first assigned each of the 76 congruent scene-object trials a value of 1 and each of the 76 incongruent scene-object trials a value of 0, before calculating the Euclidean distance between each object pair and summarizing the results in a 152 × 152 RDM (Fig 2B).

From the EEG data, we extracted brain responses to each object for the time points from -200 ms to 900 ms in intervals of 5 ms, resulting in 221 time-points. Next, we calculated the correlation distance between each object pair at each time-point. This resulted in participant-specific RDMs at each time-point that summarised the neural similarity between the objects.

In the following step, we related the congruency model RDM with the brain response RDMs using Spearman correlation resulting in an RSA time-series per participant (Fig 2C). A random effects analysis assessed the model fit of the congruency model RDM and the brain RDMs at each time-point using a t-test against zero with an alpha of 0.01. To control for multiple comparisons across time we used a cluster-mass permutation test to assign p-values to clusters of significant tests (Maris & Oostenveld, 2007). For each permutation, the sign of RSA correlation time-series between the model and brain RDM was randomly flipped for each participant, before t-tests were performed on the permuted data, and the size of the largest cluster added to the permutation distribution. Finally, the cluster p-value for clusters in the original data were defined as proportion of the 10000 permutations (plus the observed cluster mass) that was greater than or equal to the observed cluster mass.

#### Semantic model analysis

The semantic model specified the semantic-feature similarity of object concepts based on a published set of property norms (Devereaux et al., 2014). The current version of the property norms is available from the Centre of Speech, Language, and the Brain (https://cslb.psychol.cam.ac.uk/propnorms). The property norms we used summarised how 826 different concepts related to 3026 different features (e.g. a zebra ‘has stripes’, ‘eats grass’ etc), allowing us to represent each concept by a collection of features that together define the concept (e.g. a zebra ‘has legs, ‘has stripes’, but does not ‘live in trees’). We matched the objects used by Draschkow and colleagues (2018) with concepts in the property norms. A matching concept was found for 118 out of 152 objects that were presented in a congruent context, and for 116 out of 152 objects that were presented in an incongruent context. Trials containing objects for which no match could be found were excluded from further analysis. For the other trials, a semantic similarity space was defined by calculating the cosine distance between all possible pairs of objects, separately for objects that were presented in a congruent context and objects that were presented in an incongruent context, resulting in two semantic feature RDMs (Fig 2B). The resulting RDM dimensions differed across participants because while all participants viewed the same scenes, the scenes were shown to one half of the participants with a congruent object and to the other half of participants with an incongruent object. For both groups of participants, the dimension of the RDM for incongruent trials was 58 × 58, and the RDM for congruent trials for half the participants was 57 × 57 and 61 × 61 for the remaining half.

The EEG data of each participant was separated for congruent and incongruent scene-object trials, before RDMs per time-point were calculated in the same way as for the congruency model, except that now two analyses were performed, one relating the semantic feature RDM to the brain RDMs for congruent trials, and one analysis relating the semantic feature RDM to the brain RDMs for the incongruent trials (Fig 2C). Significant differences between the RSA model fit for congruent and incongruent context conditions was additionally assessed with a cluster-based permutation test using paired sample t-tests.

## Results

We combined computational modelling with RSA to test if congruency effects in N300/N400 EEG components were driven by semantic object information. First, we constructed a model of consistency to uncover when the processing of congruent and incongruent objects diverged. Then, using a model based on semantic features, we investigated the time-course of semantic processing of objects that were presented in either congruent or incongruent contexts.

### Congruency model analysis

We first assessed whether neural patterns distinguished between objects presented in congruent and incongruent environments. RSA analysis of the EEG signals revealed that the congruency model distinguished between congruent and incongruent scene-object context trials, where we saw a significant relationship between the congruency model and EEG patterns from approximately 290 to 450 ms (cluster p = 0.022; Fig 4A, Table 1). The timing of this effect is in line with previous N300/N400 effects of congruency.

**Table 1.** EEG RSA effects.

| Model RDM | time-window | cluster-mass | cluster p-value |
| --- | --- | --- | --- |
| Congruency model | 286-450 ms | 74.26 | $p = .022$ |
| Semantic model: consistent | 141-235 ms | 46.11 | $p = .064$ |
| Semantic model: inconsistent | 141-360 ms | 132.25 | $p = .005$ |
| Semantic model: inconsistent | 616-765 ms | 80.78 | $p = .021$ |
| Semantic model: incon > con | 276-395 ms | 50.25 | $p = .049$ |
| Semantic model: incon > con | 596-870 ms | 169.26 | $p = .002$ |

**Figure 3.**
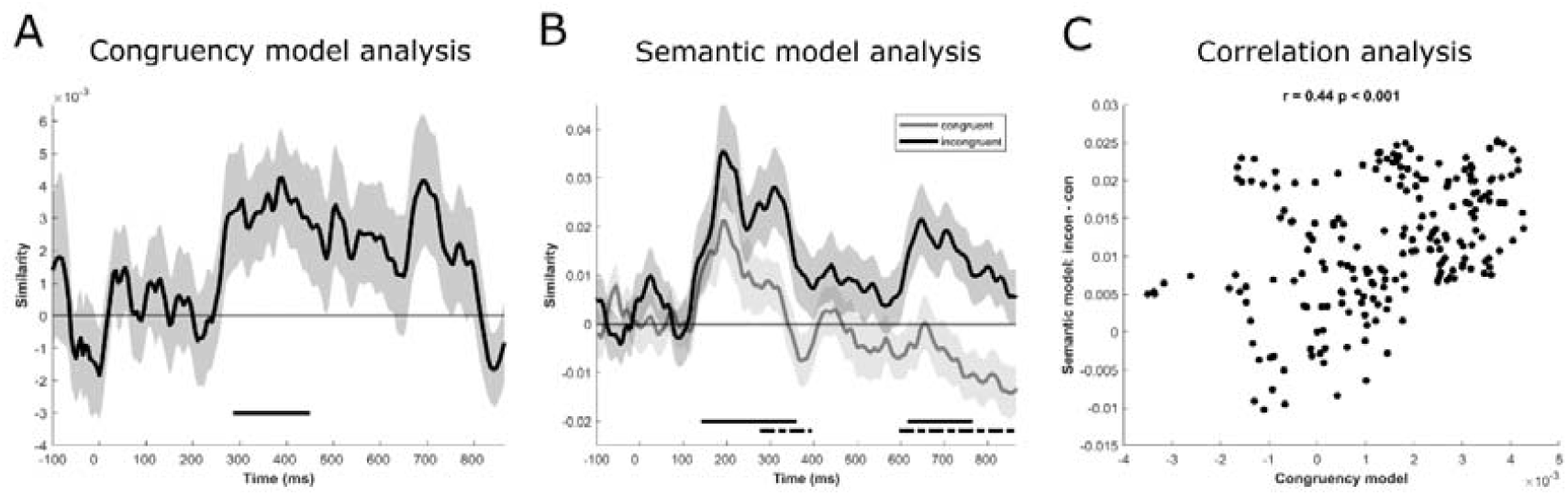
RSA results. (A) The consistency model fit shows the similarity based on Spearman correlation between the model RDM and the EEG RDMs at each time-point. Shaded area shows +-1 standard error of the mean. The horizontal bar shows a statistically significant cluster. (B) The semantic model fit is shown separately for congruent (grey line) and incongruent (black line) conditions. The horizontal bars show statistically significant clusters for incongruent objects (black solid line) and the difference of semantic model fit between congruent and incongruent objects (black dotted line). (C) Correlational analysis relating the congruency model to the difference of semantic model fit between congruent and incongruent objects.

### Semantic model analysis

While the congruency effect shows that object processing and scene information interact, it does not tell us about how semantic knowledge may be accessed differentially depending on the scene context. To address this, we constructed a semantic model based on semantic features that collectively describe each of the concepts. Two separate RDMs were created, one for objects that were presented in a congruent context and one for objects that were presented in an incongruent context. The model RDMs were then correlated with the corresponding brain RDMs, i.e. the congruent model RDM was related to brain activation RDMs to congruent objects and the incongruent model RDM to brain activation RDMs to incongruent objects.

For congruent objects, although the semantic model showed no statistically significant relationship to brain activity, it numerically performed well from around 140 ms to 235 ms (cluster p = 0.06). For incongruent objects, the semantic model fitted the brain responses significantly during two clusters, the first cluster including time-points around 140 ms to 360 ms (cluster p = 0.005) and a second cluster with time-points between around 620 to 765 ms (cluster p = 0.021; Fig 4B, Table 1). This might suggest that while semantic effects in both conditions seem to begin at similar times, approximately 150 ms after object onset, semantic effects for incongruent objects continue for a longer period of time. In order to test this, we compared the two conditions directly. The difference in model-fit between the two conditions was significant at two clusters, an early cluster including the time-points around 280 ms to 395 ms (cluster p = 0.049) and a second cluster including the time-points from approximately 600 ms to 870 ms (cluster p = 0.002; Fig 4B, Table 1). While cluster-based permutation testing does not allow precise estimation of effect on- and off-sets (Sassanhagen & Draschkow, 2019), qualitatively the time-windows of these clusters overlap with both our effects of the congruency model RDM, and known congruency effects in the N300/N400 (e.g. Draschkow et al., 2018; Ganis & Kutas, 2003; Lauer et al., 2018; Lauer et al., 2020; Mudrik et al., 2010; Mudrik et al., 2014; Võ & Wolfe, 2013), in addition to a regularly reported later effect coinciding with the P600 (e.g. Ganis & Kutas, 2003; Mudrik et al., 2010; Sauvé et al., 2017; Võ & Wolfe, 2013).

### Correlation analysis

The model-based analysis revealed overlap between the congruency and the semantic model RDMs in the time-window from approximately 290 to 395 ms. Within this time-window, the congruency model successfully distinguished if an object was presented in congruent or incongruent context, and the semantic model displayed significantly better model fit with objects that were presented in an incongruent context compared to objects that were presented in a congruent context. In order to test if these two effects were related, we correlated the time courses of the congruency model fit with the time-course of the difference between the semantic model fit for the congruent and incongruent conditions. The results confirm a significant correlation (r = 0.44, p < 0.001; Fig 4C), demonstrating that the two analyses could be capturing the same temporal effect of congruency, which might suggest that effects of congruency are explained by differences in the processing of semantic object features.

## Discussion

In the current study, we directly tested if scene context influenced object recognition through the modulation of processing semantic object information. We related the similarity based on EEG activity in response to visual objects with both a model of congruency and a semantic model that was based on semantic object property norms. Both the congruency model and semantic model captured an effect of scene context on object processing in the time-window for which N300/N400 effects were previously reported (e.g. Draschkow et al., 2018; Ganis & Kutas, 2003; Lauer et al., 2018; Lauer et al., 2020; Mudrik et al., 2010; Mudrik et al., 2014; Võ & Wolfe, 2013). Additionally, the semantic model revealed a difference in processing of congruent and incongruent objects in a later time-window beyond ∼ 600 ms, which has been reported in some previous studies (e.g. Ganis & Kutas, 2003; Mudrik et al., 2010; Sauvé et al, 2017. Võ & Wolfe, 2013). In these two time-windows, the semantic model displayed stronger fit for incongruent than for congruent objects, suggesting that the previously observed congruency effects were driven by the additional need for semantic processing of objects that were incongruent with their environment. This contrasts with alternative possibilities that semantic object information could have been accessed faster for objects in congruent compared to incongruent environments, or that semantic access was initiated at the same time and for the same duration for both congruent and incongruent conditions.

Our research is the first to employ a modelling-based approach to directly test the hypothesis that scene context influences object recognition by modulating the processing of semantic object information. The stronger model fit for incongruent objects beyond about 275 ms suggests that while both congruent and incongruent objects involve semantic processing beyond ∼150 ms, semantic processes are extended in the incongruent condition. It may well be that this extended semantic processing for incongruent objects is what underpins the congruency effect, which seems to begin at a similar time to the divergence of semantic model fits across the two conditions.

Overall, our findings are in agreement with a framework whereby context generates expectations about objects we might encounter, and thereby affects the way objects are processed (Bar, 2004; Oliva & Torralba, 2007; Federmeier et al., 2016; Clarke, 2019; Lauer & Võ, 2022). Our results, together with previous findings of congruency effects on N300/N400 EEG components, show stronger effects for objects that were unexpected compared to objects that were expected. This phenomenon is consistent with the predictive coding account (Friston, 2005) which states that the brain constructs prior expectations about upcoming sensory events based on experience, and generates an error response if the event does not match the expectation. In terms of scene-object congruency, exposure to a scene context could create a prediction about what objects are likely to appear in that scene. This is even more so here, as a fixation dot appeared prior to the object indicating the location the item would appear, thus limiting the range of likely object candidates. If the object is not congruent with the scene, and hence does not fit the prediction, it triggers a prediction error response causing delayed or enhancement of brain activity that is related to object processing, like is seen for N300/N400 EEG components (e.g. Draschkow et al., 2018; Ganis & Kutas, 2003; Lauer et al., 2018; Lauer et al., 2020; Mudrik et al., 2010; Mudrik et al., 2014; Võ & Wolfe, 2013; for a review see Lauer & Võ, 2022). Similar effects of context on object recognition have been reported not only for scenes but also for other types of prior information like the presence of other objects (Auckland et al., 2007; Kovalenko et al., 2012; McPherson & Holcomb, 1999), and has been demonstrated in semantic priming studies (Renoult et al., 2012). This indicates that the semantic effects we see here reflect a more general mechanism that is not restricted to scenes, whereby the context activates semantic or schema-consistent information within which the semantics of a new item are to be integrated, thus allowing us to use semantic information from the world around us to predict what to expect - be this what objects are likely to appear around the corner, or what words might be next in a sentence (for reviews see Kutas & Federmeier, 2011; Federmeier et al., 2016; Võ et al., 2019; Võ, 2021). Taken together, having a prior expectation about likely objects might allow for more efficient semantic processing, with the consequence that we see rapid and short semantic effects for congruent objects, and extended semantic effects for incongruent objects.

In their original work, Draschkow and colleagues (2018) demonstrated how congruency effects influencing the N300 and N400 components constitute highly related processes which allow the decoding of congruency across the two time-windows, finding significant cross-decoding of congruency from about 200 ms after object onset. The consistency model analysis that we employed, likewise tested to distinguish between objects that were presented either in congruent or incongruent context. Our results highlight a similar time-window like the decoding analysis, thus confirming that a model-based approach is suitable to capture congruency effects in EEG data.

In addition to the congruency effects in the N300/N400 time-window, the semantic model analysis revealed a difference in processing of congruent and incongruent objects in a later time-window beyond ∼600 ms. Effects in this time window were previously reported in similar studies, in which objects were embedded into scenes (e.g. Ganis & Kutas, 2003; Mudrik et al., 2010; Sauvé et al., 2017; Võ & Wolfe, 2013). However, the exact nature of these later effects remains unclear as they vary depending on task-demands (Ganis & Kutas, 2003; Võ & Wolfe, 2013; Sauvé et al., 2017). Ganis and Kutas (2003), for example, reported two different effects in this time window - a stronger positivity for incongruent objects when the task was to identify the object, and a topographically distinct effect that was stronger for congruent objects when participants were additionally instructed to provide confidence ratings. In our study, participants were required to report if a scene-object combination had been shown previously, and we find a stronger representation of semantic information when the object was incongruent, consistent with the first effect found by Ganis & Kutas (2003). As such, it is unlikely that our late effect reflects the confidence of a decision-making process. Võ and Wolfe (2013) reported a P600 component specifically in the context of mild syntactic violations of an object’s position in scenes (object misplaced), but not for extreme syntactic violations (object in impossible position, for example in the air). One possible explanation for finding these effects in the current data is that objects were embedded in scenes, and this has likely induced a combination of semantic and mild syntactic violations in incongruent scene-object trials. For example, if a ball is embedded in a photo of a kitchen and is placed inside a microwave, then in addition to the semantic incongruency, a mild syntactic violation is also created. This combination of semantic and syntactic violations might explain RSA effects of semantic object processing in incongruent scenes during a similar time window to the previously reported P600 congruency effects.

In conclusion, our results revealed that while semantic processing begins around 150 ms after the object appears, the modulatory effect of the prior scene context starts around the onset of the N300 components, resulting in longer processing of objects that are incongruent with a scene compared to objects that are congruent. Additionally, we replicated effects in previously reported time-windows of the N300/N400 and P600 EEG components using a computational modelling approach. Importantly, our study highlights how object recognition processes are flexibly adapted based on prior information, in this case showing the dynamics associated with accessing semantic knowledge are modulated by the prior context.

## Acknowledgements

This research was funded in whole, or in part, by the Wellcome Trust [Grant number 211200/Z/18/Z to AC]. The Wellcome Centre for Integrative Neuroimaging is supported by core funding from the Wellcome Trust (203139/Z/16/Z), and M L-H Võ supported by the Hessisches Ministerium für Wissenschaft und Kunst (HMWK; project “The Adaptive Mind”). For the purpose of open access, the author has applied a CC BY public copyright licence to any Author Accepted Manuscript version arising from this submission.

